# Genomic evaluation of circulating proteins for drug target characterisation and precision medicine

**DOI:** 10.1101/2020.04.03.023804

**Authors:** Lasse Folkersen, Stefan Gustafsson, Qin Wang, Daniel Hvidberg Hansen, Åsa K Hedman, Andrew Schork, Karen Page, Daria V Zhernakova, Yang Wu, James Peters, Niclas Ericsson, Sarah E Bergen, Thibaud Boutin, Andrew D Bretherick, Stefan Enroth, Anettne Kalnapenkis, Jesper R Gådin, Bianca Suur, Yan Chen, Ljubica Matic, Jeremy D Gale, Julie Lee, Weidong Zhang, Amira Quazi, Mika Ala-Korpela, Seung Hoan Choi, Annique Claringbould, John Danesh, George Davey-Smith, Federico de Masi, Sölve Elmståhl, Gunnar Engström, Eric Fauman, Celine Fernandez, Lude Franke, Paul Franks, Vilmantas Giedraitis, Chris Haley, Anders Hamsten, Andres Ingason, Åsa Johansson, Peter K Joshi, Lars Lind, Cecilia M. Lindgren, Steven Lubitz, Tom Palmer, Erin Macdonald-Dunlop, Martin Magnusson, Olle Melander, Karl Michaelsson, Andrew P. Morris, Reedik Mägi, Michael Nagle, Peter M Nilsson, Jan Nilsson, Marju Orho-Melander, Ozren Polasek, Bram Prins, Erik Pålsson, Ting Qi, Marketa Sjögren, Johan Sundström, Praveen Surendran, Urmo Võsa, Thomas Werge, Rasmus Wernersson, Harm-Jan Westra, Jian Yang, Alexandra Zhernakova, Johan Ärnlöv, Jingyuan Fu, Gustav Smith, Tonu Esko, Caroline Hayward, Ulf Gyllensten, Mikael Landen, Agneta Siegbahn, Jim F Wilson, Lars Wallentin, Adam S Butterworth, Michael V Holmes, Erik Ingelsson, Anders Mälarstig

## Abstract

Circulating proteins are vital in human health and disease and are frequently used as biomarkers for clinical decision-making or as targets for pharmacological intervention. By mapping and replicating protein quantitative trait loci (pQTL) for 90 cardiovascular proteins in over 30,000 individuals, we identified 467 pQTLs for 85 proteins. The pQTLs were used in combination with other sources of information to evaluate known drug targets, and suggest new target candidates or repositioning opportunities, underpinned by a) causality assessment using Mendelian randomization, b) pathway mapping using *trans*-pQTL gene assignments, and c) protein-centric polygenic risk scores enabling matching of plausible target mechanisms to sub-groups of individuals enabling precision medicine.

## Main

Proteins circulating in blood are derived from multiple organs and cell types, and consist of both actively secreted and passively leaked proteins. Plasma proteins are frequently used as biomarkers to diagnose and predict disease and have been of key importance for clinical practice and drug development for many decades.

Circulating proteins are attractive as potential drug targets as they can often be directly perturbed using conventional small molecules or biologics such as monoclonal antibodies^1^. However, a prerequisite for successful drug development is efficacy, which is predicated on the drug target playing a causal role in disease. One approach to clarifying causation is through Mendelian randomization (MR), which has successfully predicted the outcome of randomized controlled trials (RCT) for pharmacological targets such as PCSK9, LpPLA2 and NPC1L1, and is increasingly becoming a standard tool for triaging new drug targets^2^.

Recent technological developments of targeted proteomic methods have enabled hundreds to thousands of circulating proteins to be measured simultaneously in large studies^3,4^. This has paved the way for studies of genetic regulation of circulating proteins using genome-wide association studies (GWAS) for detection of protein quantitative trait loci (pQTL)^3–5^.

Here, we present a genome-wide meta-analysis of 90 cardiovascular-related proteins, many of which are established prognostic biomarkers or drug targets, measured using the Olink Proximity Extension Assay CVD-I panel ^6^ in 30,931 subjects across 14 studies. The identified pQTLs were combined with other sources of information to suggest new target candidates underpinned by insights into *cis*- and *trans*-regulation of protein levels and to evaluate past and present efforts to therapeutically modify the proteins analysed in the present investigation. We also show that protein-centric polygenic risk scores (PRS) can predict a substantial fraction of inter-individual variability in circulating protein levels, explaining a proportion of disease susceptibility attributable to specific biological pathways.

These are the first results to emerge from the SCALLOP consortium, a collaborative framework for pQTL mapping and biomarker analysis of proteins on the Olink platform (www.scallop-consortium.com).

## Results

### Genome-wide meta-analysis of 90 proteins in 21,758 human subjects across 13 studies reveals 467 independent genetic loci associated with plasma levels of 85 proteins

Ninety proteins in up to 21,758 participants from 13 cohorts passed quality control (QC) criteria and were available for GWAS meta-analysis [Supplementary Table 1]. In addition to standard conventions, we used between-study heterogeneity to guide our P-value threshold used to denote GWAS significance. In the presence of between-study heterogeneity (*P-het*<9×10^−5^), SNPs had to surpass a discovery GWAS threshold that took all 90 proteins tested into account (*P*<5.6×10^−10^) and replicate at a nominal P-value threshold (P<0.05) in two separate studies (9,173 individuals) with directionally concordant beta coefficients for us to call the pQTL. In the absence of between-study heterogeneity, in order to avoid false negatives, we relaxed our *P*-value threshold for discovery to that of conventional GWAS (*P*<5×10^−8^) in a meta-analysis of the discovery and replication datasets. Using these criteria, 344 uncorrelated (r^2^=0) SNPs (75 *cis*-and 269 *trans*-pQTL) showed association with 85 proteins [Figure 1] [Supplementary Table 2]. Fifty-seven additional SNPs that did not fulfil the above metrics, but surpassed conventional GWAS thresholds in the discovery stage (*P*<5×10^−8^) are also presented as suggestive [Supplementary Figure 1]. Conditioning on each of the pQTLs using the GCTA-COJO software, we identified an additional 123 secondary pQTLs meeting our GWAS thresholds as defined above, and 21 suggestive secondary pQTLs that surpassed conventional genome-wide significance [Supplementary Table 2]. Some proteins such as SCF, RAGE, PAPPA, CTSL1 and MPO showed association with more than ten primary pQTLs, but most proteins (22 of 85) were associated with 2 primary pQTLs. We also observed that some proteins were associated with multiple conditionally significant (secondary) pQTLs such as CCL-4 with 4 secondary signals, implicating complex genetic regulation of circulating CCL-4 at the *CCL4* locus.

**Figure 1.**
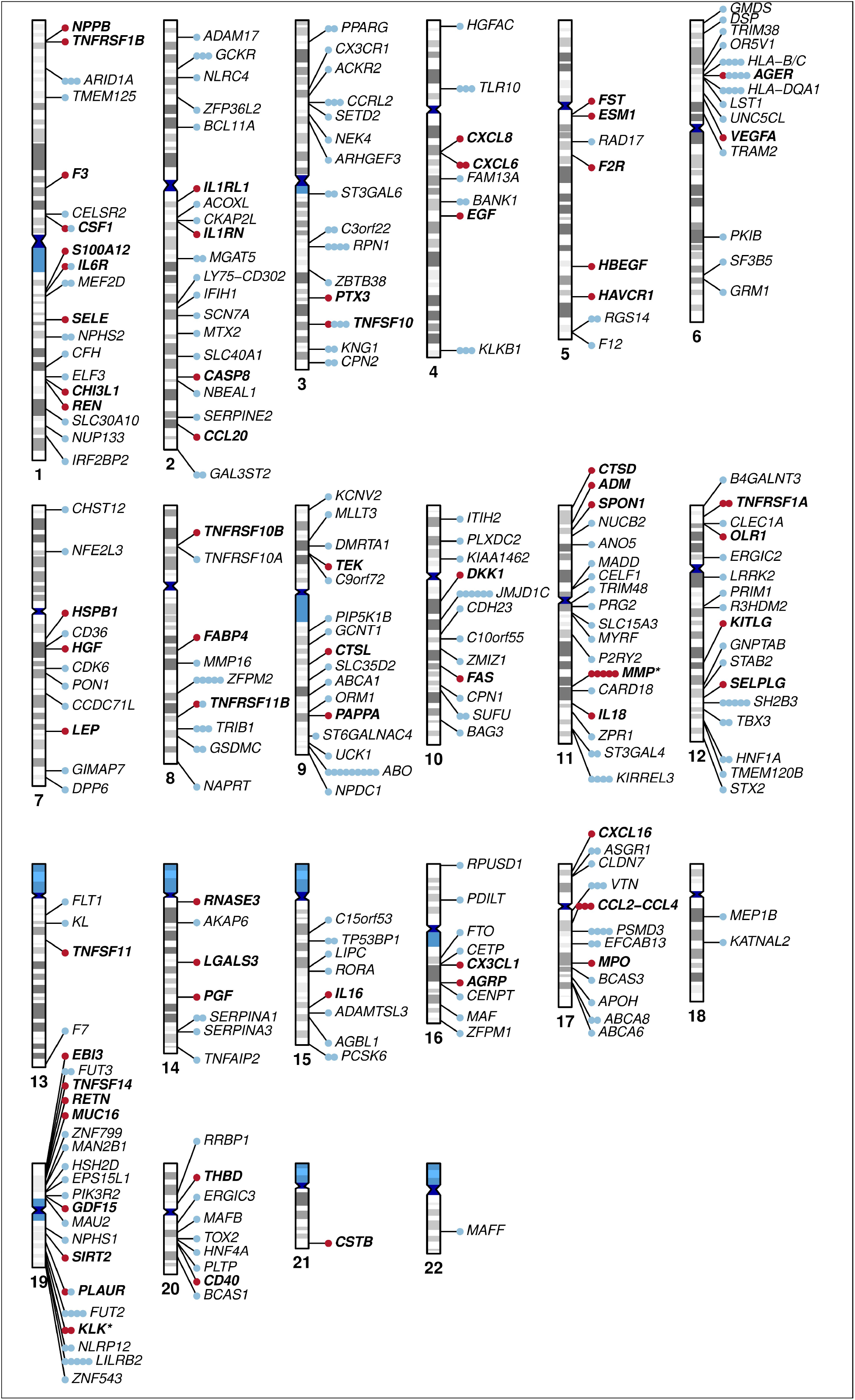
Chromosomal location of all primary associations passing GWAS significance, here defined as variants surpassing P<5.6×10^−10^ with replication at nominal P<0.05, or for non-heterogeneous variants (P<9×10^−5^), surpassing a conventional GWAS threshold of P<5×10^−8^ in the joint discovery and replication meta-analysis. *Cis*-pQTLs are shown in red (bold) and trans-pQTLs in blue. The gene annotations refer to the gene closest to the pQTL.

### Analysis of *trans*-pQTLs suggests that transcriptional regulation, post-translational modification, cell-signalling and protease activity are common mechanisms by which genetic variants affect plasma protein levels

A “best guess” causal gene for each of the CVD-I trans-pQTLs was assigned by a hierarchical approach based on analysis of protein-protein interactions (PPI), literature mining [Supplementary Table 3], genomic distance to gene and manual literature review. In total, 239 primary significant trans-pQTLs were assigned to unique genes and 30 trans-pQTLs were assigned more than one gene, with *ABO, ST3GAL4, JMJD1C, SH2B3, ZFPM2* showing association with the levels of five or more CVD-I proteins [Supplementary Figure 2B] [Supplementary Table 2]. Extending this analysis to pQTLs from literature expanded the list of genes with five or more protein associations to include also *KLKB1, GCKR, FUT2, TRIB1, SORT1 and F12* [Supplementary Table 4].

Gene ontology (GO) analysis of genes assigned to all significant trans-pQTLs showed functional enrichment for chemokine binding, glycosaminoglycan binding, receptor binding and G-protein coupled chemoattractant activity [Figure 2C]. A broader classification of genes assigned to both cis- and trans-pQTLs [Figure 2A, 2B] using a wider set of tools (Online Methods) suggested that transcriptional regulation, post-translational modifications, such as glycation and sialylation, cell-signalling events, protease activity and receptor binding are potential common mechanisms by which trans-pQTLs influence circulating protein levels. The default gene calls and paths for the CVD-I *trans*-pQTLs based on PPI and literature mining can be visualised using the <u>SCALLOP CVD-I network tool</u> [Supplementary Figure 2B] whereas details on the classification of genes are available in the Online Methods.

**Figure 2.**
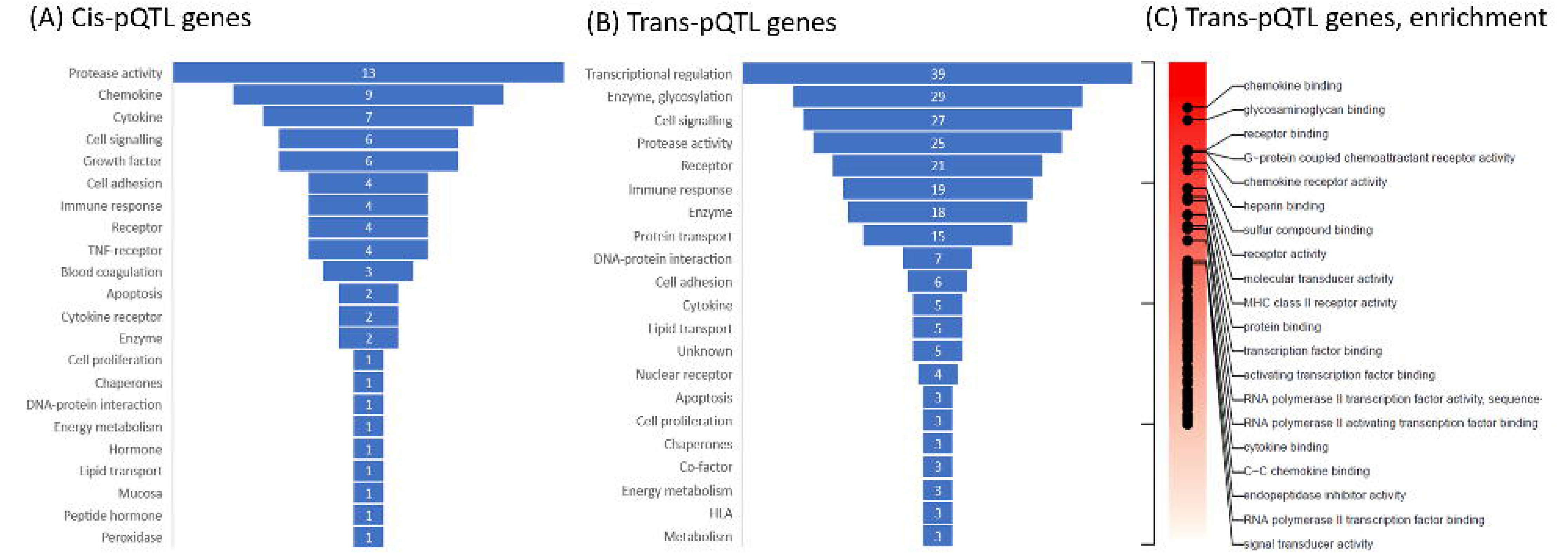
Classification of cis- and trans-pQTL genes. **A.** The gene ontology label of all cis-pQTL genes, i.e. the protein-encoding genes. **B.** The gene-ontology label of all best-guess trans-pQTL genes. **C.** Gene set enrichment analysis of genes assigned to all significant trans-pQTLs, showing the top-gene sets from the Gene Ontology set Molecular Function.

### Evidence of mRNA expression mediating associations with a third of cis pQTLs

We investigated the overlap of the CVD-I *cis*- and *trans*-pQTLs with expression quantitative trait loci (eQTL) by a combination of approaches and eQTL studies, including direct genetic lookups and co-localisation using PrediXcan ^7^ and SMR / HEIDI ^8^. For direct lookups, three studies were used: LifeLines-DEEP (whole blood), eQTLGen meta-analysis (whole blood and PBMCs) and GTEx (48 tissue types). Of 545 significant and suggestive pQTLs, eQTL data were available for 434 SNP-transcript pairs, including 168 *cis*-pQTLs and 266 *trans*-pQTLs. Of these, 72 (43%) of *cis*-pQTLs had at least one corresponding eQTL (FDR<0.05) in any of the eQTL datasets investigated, implicating 42 of the 75 proteins with a *cis*-pQTL. At a more stringent eQTL p-value of *P*<5×10^−8^, the percentage with a corresponding eQTL was 26 %, similar to some previous reports ^9–11^ [Supplementary Table 5].

Co-localisation analysis of CVD-I cis-pQTLs and mRNA levels was performed in selected tissues from the GTEx project by first imputing mRNA expression of the CVD-I protein-encoding transcripts using the PrediXcan^7^ algorithm in one of the SCALLOP CVD-I cohorts (IMPROVE), and then testing imputed mRNA levels for association with CVD-I plasma protein levels using linear regression. Twenty-six of the 90 CVD-I proteins were associated with their corresponding mRNA transcript (FDR<0.05) in at least one of the 20 GTEx tissues investigated [Supplementary Figure 3]. All 26 proteins were among the 42 proteins found to also be an eQTL by direct lookups. Proteins CCL4, CD40, CHI3L1, CSTB and IL-6RA all associated with their corresponding transcript across five or more tissues whereas proteins ST2 and RAGE showed significant association exclusively in lung, and CTSD exclusively in skeletal muscle.

Next, we used the SMR and HEIDI methods^8^ to test for pleiotropic associations between plasma protein and mRNA expression (note that the HEIDI method attempts to reject association because of LD between pQTL and eQTL). In total, 125 associations between 96 genes and 54 proteins were identified at an experiment-wise SMR test significance level (*P_SMR_*<0.05/8558) and a stringent HEIDI test threshold (P_HEIDI_ > 0.01) [Supplementary Table 6], of which 23.2 % were in *cis*-pQTL regions, such as IL-8 and U-PAR. The 96 genes were located in 74 loci, suggesting that pleiotropic associations between protein and mRNA expression were present for 18.4 % of significant and suggestive primary loci using SMR / HEIDI.

### A minor proportion of *cis*-acting pQTLs are in high linkage-disequilibrium with non-synonymous coding variants

“Pseudo-pQTLs” caused by epitope effects, i.e. differential assay recognition depending on presence of protein-altering variants, is a theoretical possibility for *cis*-pQTLs and likely dependent on the method of protein quantification ^4,12^. To evaluate the potential for pseudo-pQTLs among the CVD-I pQTLs, we investigated presence of protein-altering variants for sentinel variants or variants in high linkage disequilibrium with a sentinel variant. Of the 90 proteins, 85 had at least one pQTL, including 12 with only *cis*-pQTLs, 10 with only *trans*-pQTLs and 63 with both *cis*- and *trans*-pQTLs. Of the 170 primary or secondary *cis*-pQTLs for 75 proteins, 20 *cis*-pQTLs for 18 proteins had a sentinel variant in high linkage disequilibrium (LD; R^2^>0.9) with a protein-altering variant, which suggests potential to affect assay performance [Supplementary Table 1]. Of the 20 *cis*-pQTLs with a sentinel variant in high LD with a protein-altering variant, seven were not associated with mRNA expression in any of our analyses [Supplementary Table 5][Supplementary Table 6] [Supplementary Figure 3], indicating potential for epitope effects, requiring further validation in future studies using orthogonal assays.

### Orthogonal evidence based on pharmacological intervention and transgenic mice supports causal gene to protein relationships for a subset of the CVD-I *trans*-pQTLs

Of the 269 *trans*-pQTLs identified, eight were assigned to gene products targeted by compounds or antibodies that have been in clinical development [Supplementary Table 7]. Assuming that *trans*-pQTLs represent causal relationships between gene variants and proteins, we hypothesized that the downstream CVD-I proteins associated with CVD-I *trans*-pQTL genes would be modulated on therapeutic modification of the gene product. Support for this hypothesis was obtained by previous work showing that circulating FABP4 is upregulated upon treatment with glitazones (PPARG inhibitors)^13^; that circulating IL-6 is increased after treatment with tociluzumab^14^ (IL6R inhibitor) and that circulating TNF-R2 is decreased upon infliximab (TNFA inhibitor) treatment in patients with Crohn’s disease^15^, which supports CVD-I *trans*-pQTLs for these proteins. Along these lines, we present novel evidence supporting our *trans*-pQTL analysis implicating *CCR5* in plasma CCL-4 levels and *CCR2* in plasma MCP-1, which are targeted in combination by the small-molecule dual-inhibitor PF-04634817 ^16^. To test whether dual inhibition of CCR5 and CCR2 resulted in a change on circulating CCL-4 and MCP-1 respectively, we measured the plasma protein levels of CCL-4, MCP-1, CCL-3, CCL-5 (RANTES), CCL-8, as well as 10 additional Olink CVD-I proteins in 350 type 2 diabetes patients in a randomized, double-blind, placebo-controlled phase-II trial evaluating the efficacy of PF-04634817 in diabetic nephropathy (NCT01712061). Compared to placebo, we observed a 9.25-fold increase in circulating MCP-1 levels (p < 0.0001) and a 2.11-fold increase in circulating CCL4 levels (p< 0.0001) at week 12 [Figure 3]. An alternative ligand for CCR-2; CCL-8 did not change following exposure to PF-04634817, and neither did other CCR-5 ligands, such as CCL-5 (RANTES) and CCL-3. Moreover, EN-RAGE, FGF-23, KIM-1, myoglobin and TNFR-2 were unchanged following PF-04634817 exposure [Supplementary Figure 4]. We conclude that CVD-I *trans*-pQTLs at *CCR5* and *CCR2* were concordant with the effects of PF-04634817 in human.

**Figure 3.**
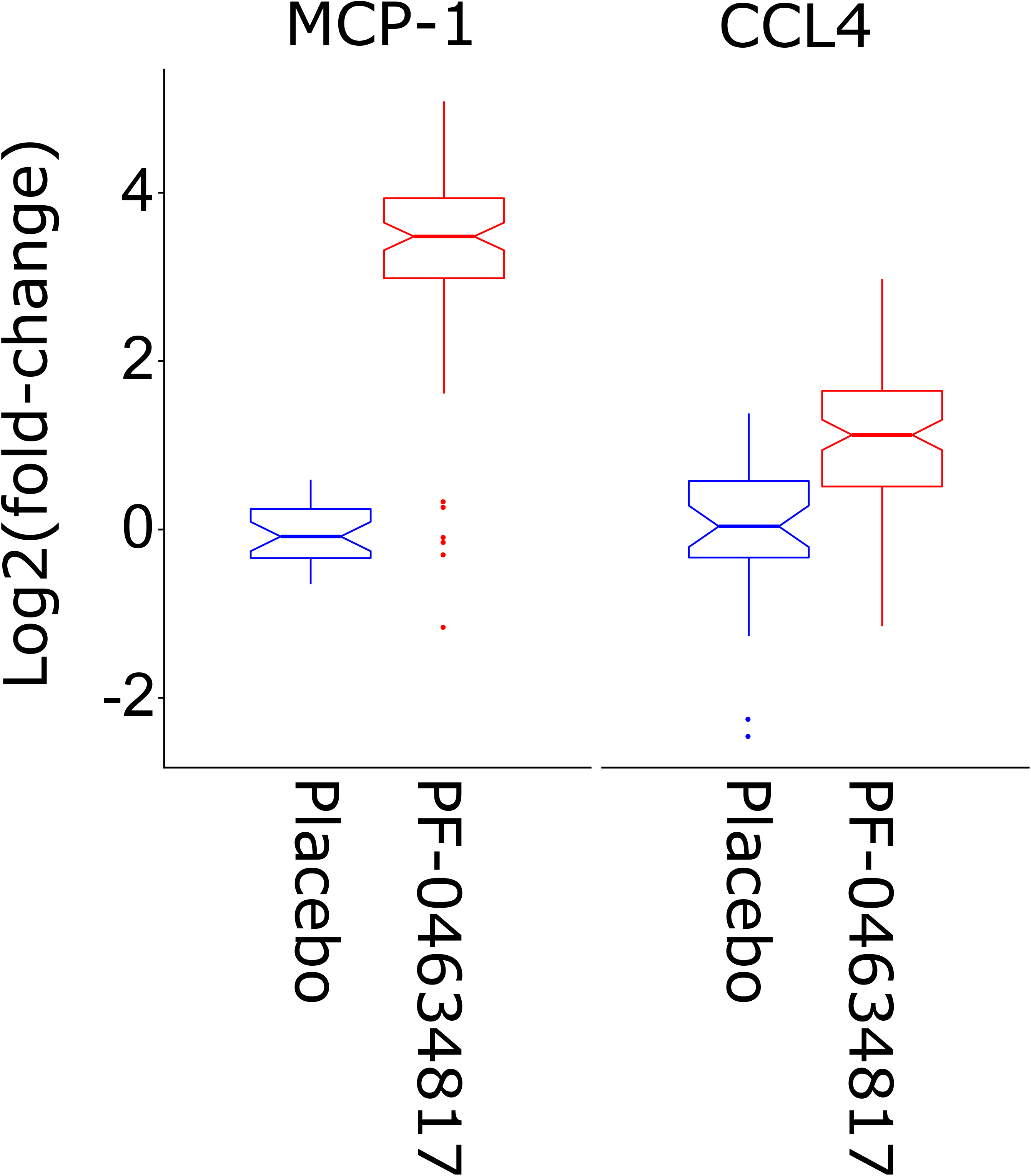
Plasma levels of MCP-1 and CCL4 in human subjects treated with a small-molecule dual-inhibitor of CCR5 and CCR2 (PF-04634817) or placebo. Induction of MCP-1 and CCL4 upon inhibition of CCR5 and CCR2 mirrors the observed CVD-I trans-pQTLs.

Two of the genes implicated by CVD-I *trans*-pQTLs, *ABCA1* and *TRIB1* for circulating SCF levels, were also investigated in the mouse. Mice with liver-specific or whole-body knockdown of *ABCA1*^17^ and *TRIB1*^18^ respectively showed decreased plasma levels of SCF compared to matched wild-type controls [Figure 4], concordant with the human CVD-I *trans*-pQTLs.

**Figure 4.**
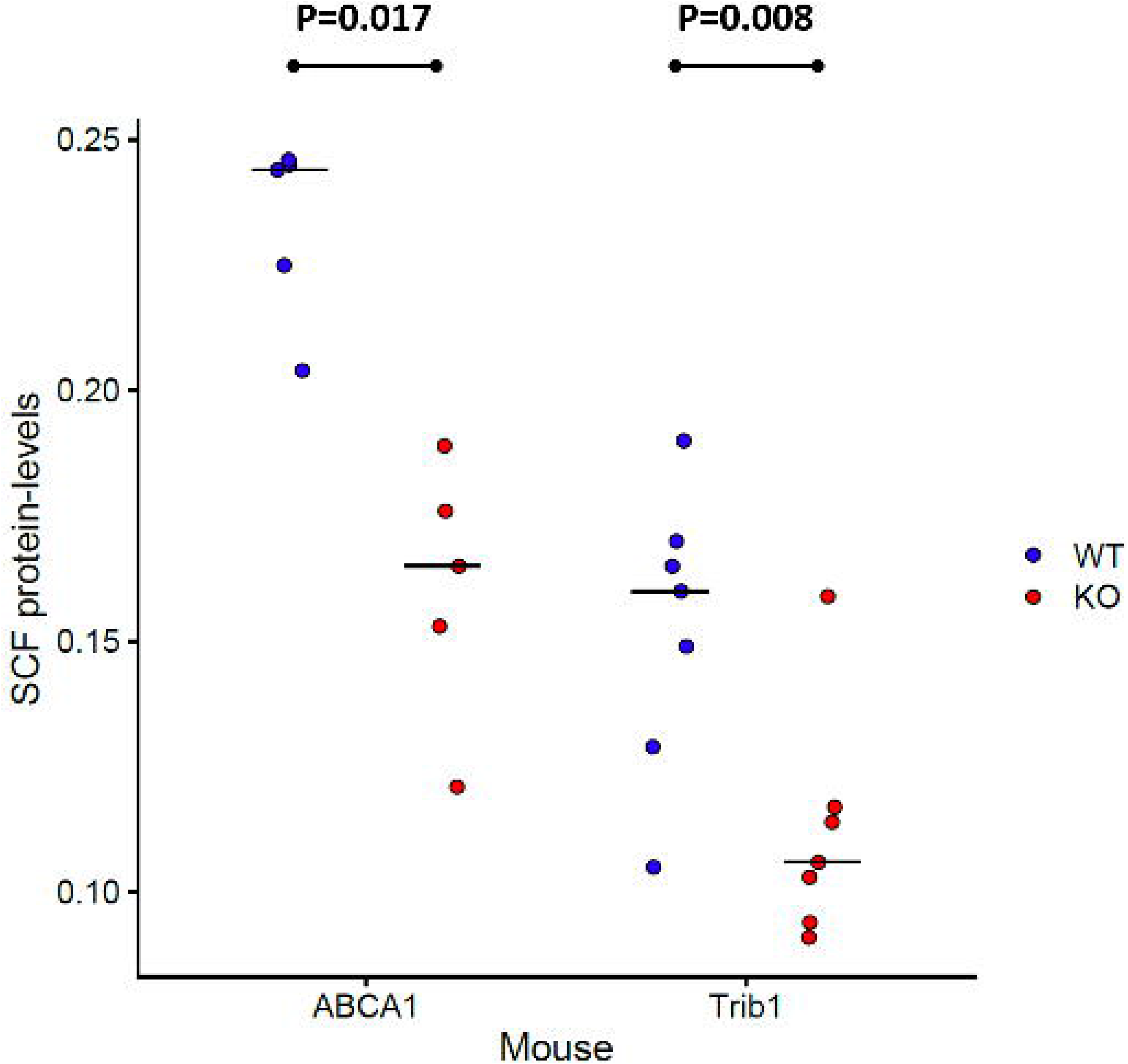
Plot showing plasma levels of SCF in *ABCA1* and *TRIB1* transgenic mice compared to wild-type controls. Knockdown of *ABCA1* or *TRIB1* resulted in decreased circulating SCF levels mirroring CVD-I trans-pQTLs for SCF. Shown in the plot are SCF levels of individual mice represented by filled circles (wild-type in blue and transgenic mice in red) and the median level per group.

### Mendelian randomization analysis revealed 25 CVD-I proteins causal for at least one human complex disease or phenotype with strong evidence. Of those, 7 proteins were concordant with launched therapies or ongoing clinical stage drug development

To identify potential causal disease pathways indexed by proteins, we conducted an MR analysis of 85 proteins across 38 outcomes. 25 proteins showed strong evidence of causality for at least one disease or phenotype and an additional 24 proteins showed intermediate evidence of causality. [Figure 5A; Supplementary Figure 5]. Using open-source information (clinicaltrials.gov) (www.ebi.ac.uk/chembl/) (www.drugbank.ca/) (www.opentargets.org) and Clarivate Integrity (integrity.clarivate.com), we identified records on past or present clinical drug development programs for 14 of the 25 proteins, all of which have been in phase 2 trials or later [Supplementary Table 7]. Of the 14 proteins, seven proteins were targeted for an indication different from the phenotype implicated by our MR analysis. Eleven of the 25 proteins have never been targeted in clinical trials, but may provide new promising target candidates for indications closely related to the traits in the MR analysis.

**Figure 5.**
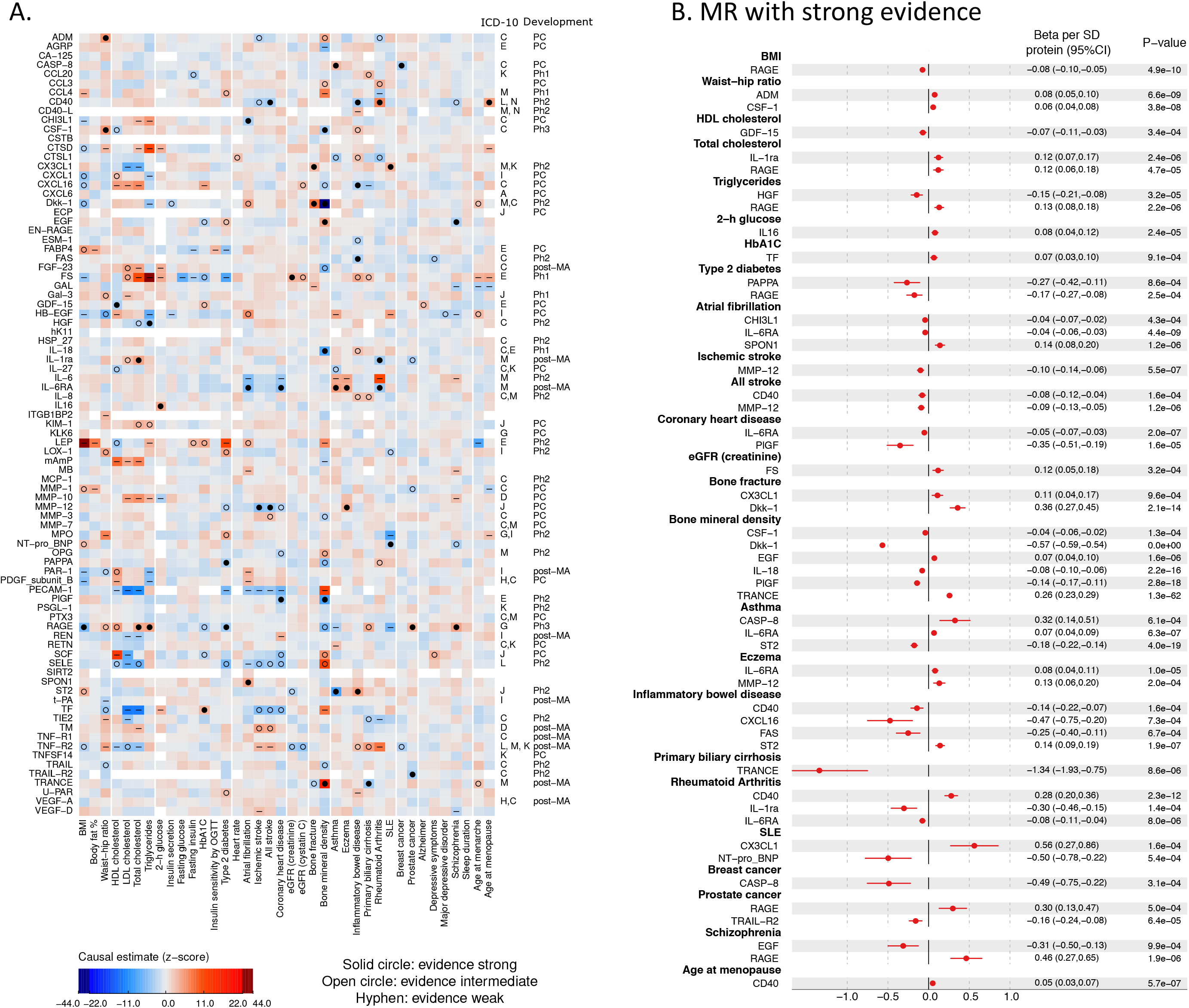
**A.** Heatmap of Mendelian randomization analyses of 38 complex traits. **B.** Forest plot showing CVD-I proteins with strong evidence of causality in the Mendelian randomization analysis.

Several published MR findings were confirmed, including that *IL6RA* variants associated with higher circulating levels of interleukin-6 (IL-6) and soluble IL6-RA were associated with lower risk of coronary heart disease (CHD), rheumatoid arthritis (RA) and atrial fibrillation but higher risks of atopy, such as asthma and eczema^19^. We also replicated previous findings suggesting a causal contribution of IL-1ra to rheumatoid arthritis (RA) but an inverse causal relationship with cholesterol levels ^20^, and a protective role of genetically higher MMP-12 against stroke ^4,21^.

Some novel MR observations included higher levels of CD40 protein and increased risk of RA, higher MMP-12 and increased risk of eczema, and higher TRAIL-R2 proteins levels and prostate cancer. Further, Dkk-1 has been targeted by a humanised monoclonal antibody (DKN-01) in clinical trials for advanced cancer (NCT01457417, NCT02375880), and was in our study causally linked to higher risk of bone fractures and lower risk of estimated bone mineral density (eBMD). In addition, strong evidence for protective roles of PLGF in CHD, CASP-8 in breast cancer and ST2 in asthma was observed. RAGE was causally linked to several traits, including lower body mass index (BMI) and a corresponding lower risk of type 2 diabetes (T2D), higher total cholesterol and triglycerides and higher risk of prostate cancer and schizophrenia. A small molecule brain penetrant RAGE inhibitor was tested in a phase 2 trial of Alzheimer’s disease (NCT00566397), but was stopped early for futility. We saw no strong signal for Alzheimer’s disease (or vascular disease) in our MR analysis. Our findings identify potential target-mediated effects across multiple other complex phenotypes that might manifest in beneficial and/or harmful effects on patients receiving RAGE-modifying therapies.

### Heritability analysis and polygenic risk scores (PRS) derived for each of the 85 proteins demonstrates large differences in genetic architecture

We calculated SNP-heritability contributed by the major reported loci (major loci h_SNP_^2^) [supplementary table 2], as well as additional genome-wide SNP-heritability (polygenic h_SNP_^2^) for each protein included in the SCALLOP CVD-I meta-analysis. We observed a large range of different genetic architectures: Differences in magnitude of the genetic component (h_SNP_^2^) ranged from 0.01 (EGF) to 0.46 (IL-6RA). Differences in the contribution from non-genome-wide significant SNPs ranged from essentially monogenic (e.g. IL-6RA) to others showing considerable locus heterogeneity with genetic contributions originating entirely from a polygenic background with no single dominating locus (e.g. PDGF-B and Galanin) [Figure 6B].

**Figure 6.**
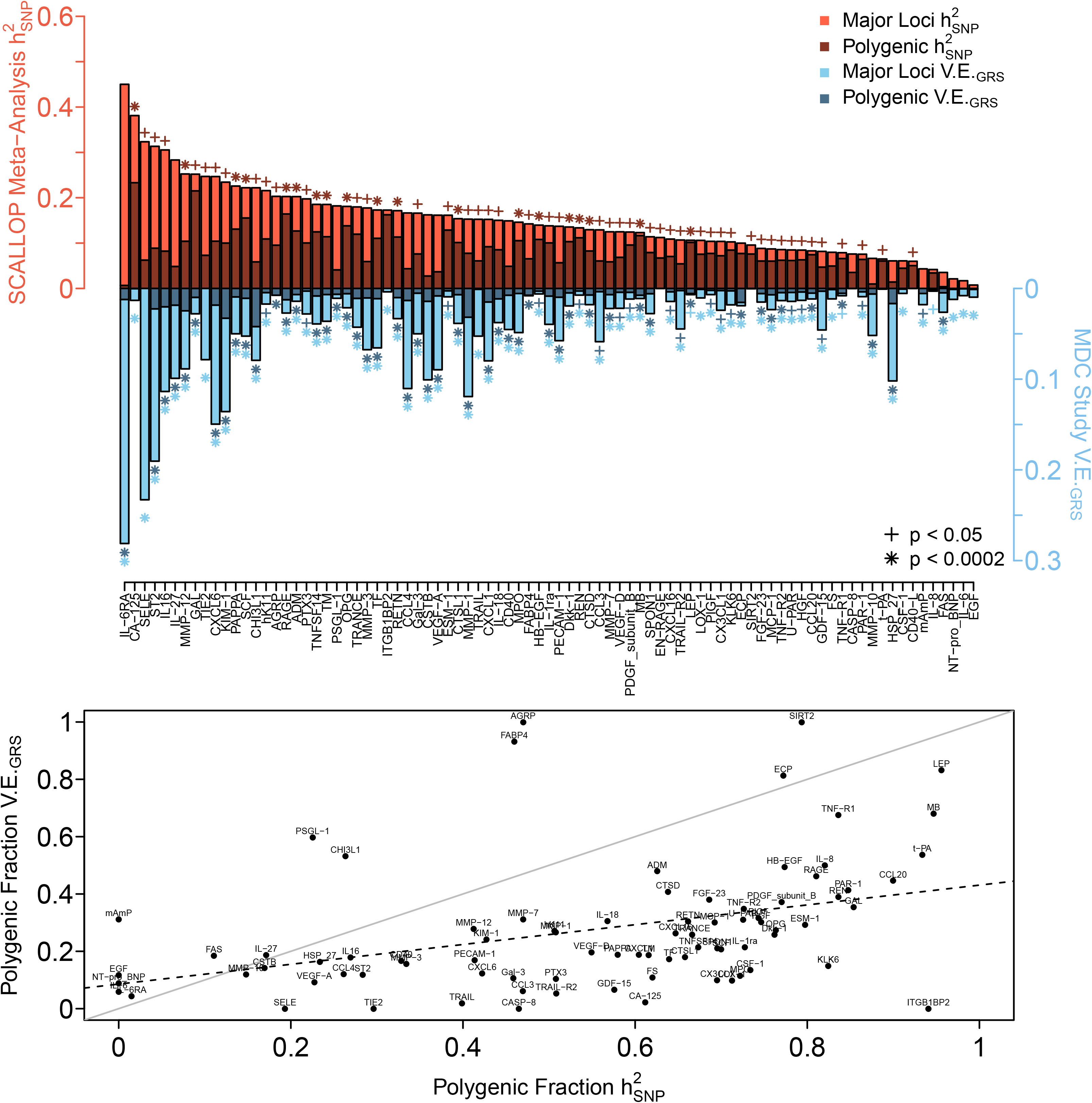
**A.** SNP-Heritability in the SCALLOP consortium discovery cohorts stratified by contributions major loci (light red) and polygenic effects (dark red). In the independent MDC cohort, additional variability explained by adding major loci (light blue) and polygenic risk scores (dark blue). **B.** Differences in how protein levels are affected by polygenic (non-genome-wide significant) loci vs major loci, shown for both the SCALLOP consortium discovery cohorts as h_SNP_^2^ and for the MDC cohort as variability explained.

In addition, we calculated the out of sample variance explained in the independent Malmo Diet and Cancer (MDC) study (N~4,500) both for genome-wide significant loci (major loci V.E._PRS_), as well as additional variance explained by adding PRS (polygenic V.E._PRS_) [Figure 6A]. The protein PRS’ applied in the MDC study for 11 proteins exceeded 10 % of variance explained (V.E._PRS_) and the PRS’ for another 14 proteins exceeded 5 % of variance explained, suggesting that the genetic contribution to inter-individual variability of CVD-I protein levels is considerable.

### A polygenic risk score for circulating ST2 levels shows a dose-response relationship with asthma

Since circulating ST2 showed strong evidence of causation in asthma and inflammatory bowel disease (IBD) and the polygenic V.E._PRS_ model for ST2 explained nearly 20 % of its variance, we attempted to quantify the effect of the ST2 polygenic V.E._PRS_ on circulating ST2 levels in the MDC study, and risk of asthma and IBD in 337,484 unrelated White British subjects in the UK Biobank. The range of circulating ST2 across 11 categories of the ST2 PRS in MDC was nearly 1.2 standard deviations [Figure 7A]. Corroborating the Mendelian randomization analysis, the ST2 PRS showed a strong negative dose-response relationship with risk of asthma (p=1.2×10^−8^) and a positive trend for risk of IBD (p=0.13) [Figure 7B and C]. Overlaying the linear trends for ST2 levels, asthma and IBD using meta-regression, an increase in the PRS equivalent to a 1 standard deviation higher circulating ST2, corresponded to a 8.6 % (95%CI 3.8%, 13.2%; P=0.004) reduction in the relative risk of asthma and a 4.3 % (95%CI-3.8%, 13.0%; P=0.263) increase in the relative risk of IBD [Supplementary Figure 8].

**Figure 7.**
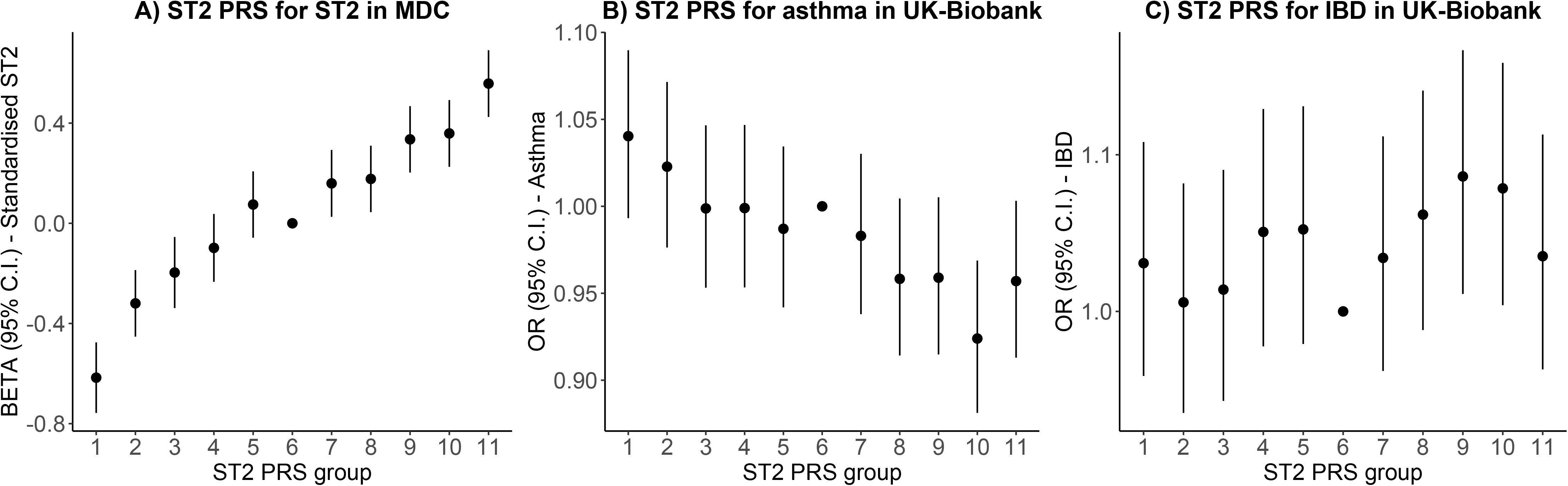
**A.** Association of a polygenic risk score (PRS) with ST2 levels in the independent MDC cohort.. **B.** Association of the ST2 PRS with asthma in the UK-biobank. **B.** Association of the ST2 PRS with inflammatory bowel disease (IBD) in the UK-biobank. The ST2 PRS was divided into 11 quantiles, with the middle group (quantile number 6) as the reference category. Effect estimates are presented as quantile-specific mean differences (ST2) and odds ratios (asthma and IBD) relative to the reference category.

### Reverse Mendelian randomization identifies widespread causal relationships, with each of the 37 complex phenotypes affecting at least one of the CVD-I proteins

To investigate whether genetic susceptibility (liability) to complex disease and phenotypes causally alter circulating levels of CVD-I proteins, we also performed MR using 38 complex phenotypes (including continuous risk factors, such as adiposity and clinical outcomes, such as T2D) as exposure and CVD-I protein levels as outcomes. All CVD-I proteins were causally altered by at least one complex phenotype. BMI and estimated glomerular filtration rate (eGFR) causally affected 32 and 29 of the 85 tested proteins respectively [Figure 8A; Supplementary Figure 7]. BMI seemed to causally affect protein levels in both positive and negative directions, whereas only REN (renin) was causally decreased with genetically higher eGFR. In an effort to elucidate whether these estimates were recapitulated in simple observational analyses, we compared effect estimates from linear regression analyses of associations of BMI and eGFR with each respective CVD-I protein in one of the participating study cohorts (IMPROVE). The correlation between the observational and MR estimates were high for BMI (R=0.78), and more modest for eGFR (R=0.50) [Figure 8B-C].

**Figure 8.**
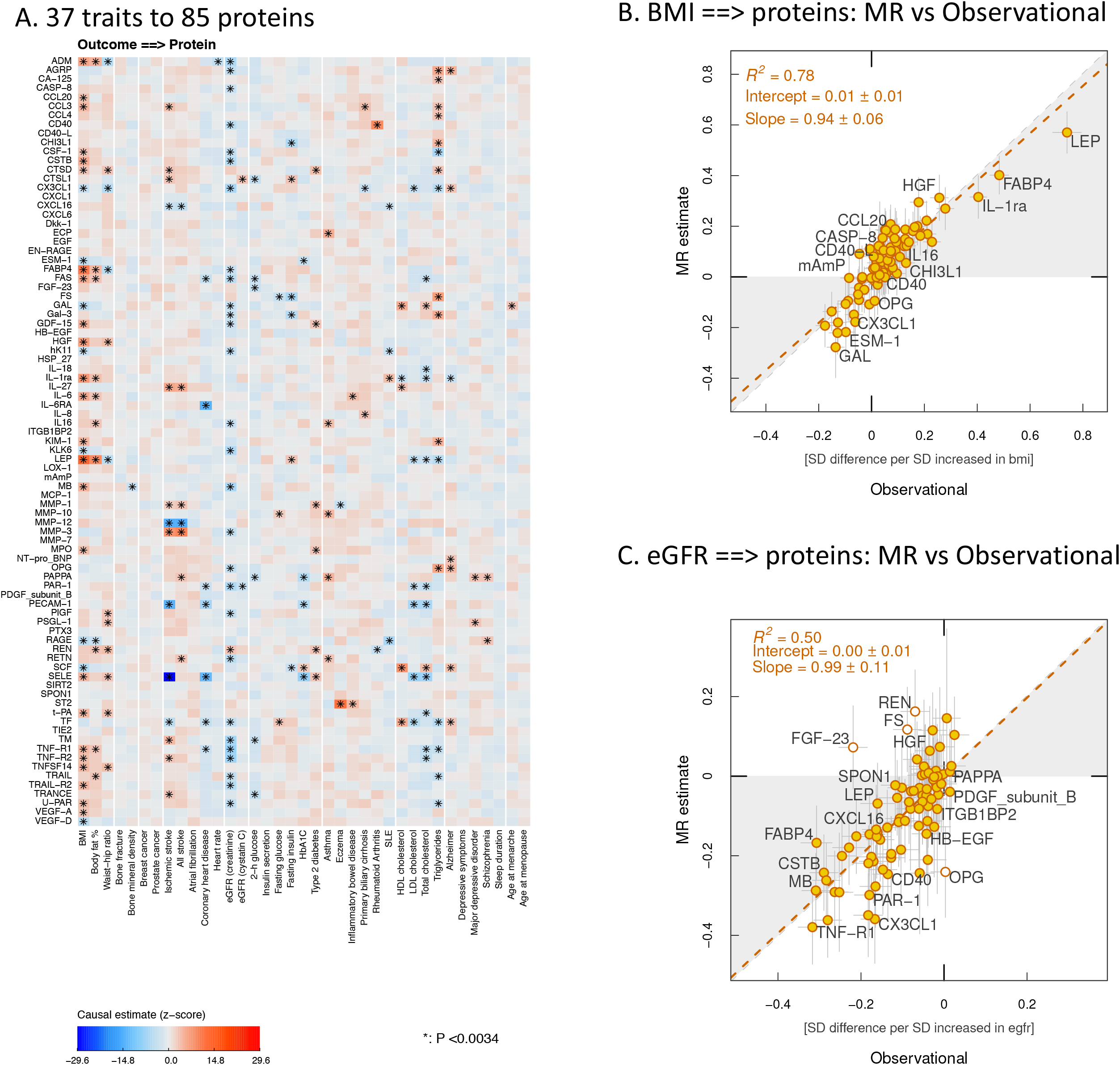
**A.** Heatmap showing the causal estimates of 38 complex traits on CVD-I protein levels. **B.** Correlation between beta-values for association between body mass index and circulating levels of CVD-I proteins in the IMPROVE cohort, and causal estimates from the Mendelian randomization analysis of body mass index genetic liability on same CVD-I proteins. **C.** Same as B but for estimated glomerular filtration rate.

## Discussion

Using a meta-analysis approach including >30,000 individuals, we identified and replicated 344 primary and 123 secondary pQTLs for 85 circulating proteins which were combined with multiple types of molecular- and genetic data to yield new insights for translational studies and drug development. Our study demonstrates that pQTLs can be harnessed to enhance evaluation of therapeutic hypotheses for protein targets, and to support those hypotheses with basic insights into potential protein regulatory pathways and biomarker strategies. However, we also observed large differences between proteins in relation to genetic architecture, suggesting that the relative strength to apply these strategies is likely protein-dependent.

Our pQTL-based framework was developed to address several key challenges associated with drug development, including a) mapping of potential regulatory pathways for circulating proteins, b) identification of new target candidates based on causal proteins, c) repositioning of drugs in development, d) target-associated safety and e) matching of target mechanisms to patients by protein biomarkers or genetic PRS’ [Figure 9].

**Figure 9.**
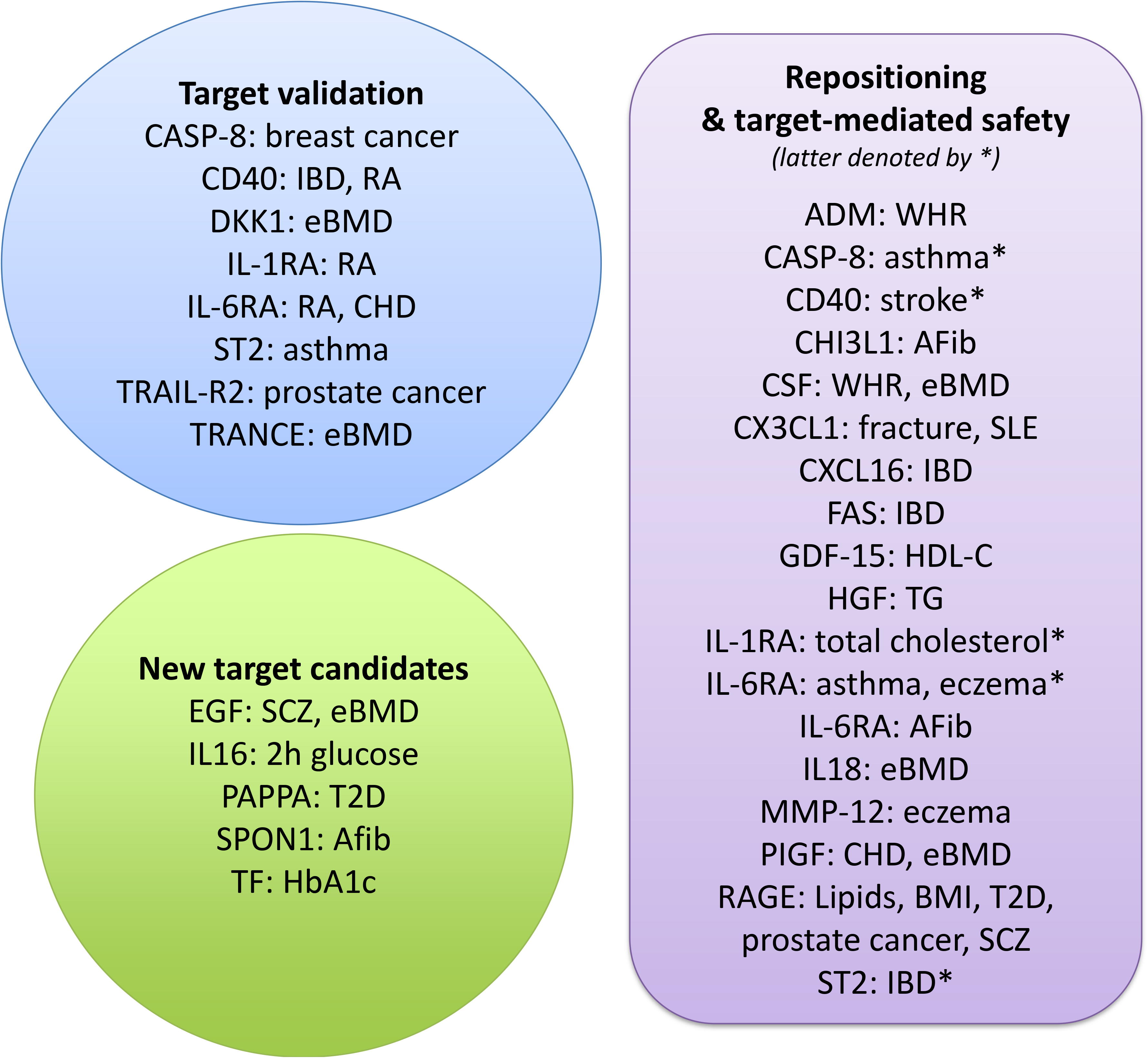
Protein-trait relationships that support target validation, repositioning, target-mediated safety and new candidates for drug development. For more information, see data presented in Supplementary Table 7.

The mapping of *trans*-pQTLs, which typically have smaller effects on protein levels, was aided by the large SCALLOP discovery sample size, yielding on average 4 independent pQTLs per protein. A causal gene was assigned for each *trans*-pQTL to generate hypotheses that can be further tested using *in vitro* or *in vivo* perturbation experiments. The robustness of causal gene assignments for a few selected *trans*-pQTLs was demonstrated using samples from a randomised controlled trial testing a dual small-molecular inhibitor of the protein products of assigned genes (*CCR5*, *CCR2*) and transgenic mice with liver-specific knockdown of assigned genes (*ABCA1*, *TRIB1*). Although further studies will be needed for orthogonal validation of most of the genes assigned from the CVD-I *trans*-pQTLs, several of the implicated genes have previously been identified as regulators of some of the CVD-I proteins including *CASP1*^22^, *NLRC4*^22^ and *GSDMD*^23^ for IL-18, *FLT1*^24^ for PLGF, *ADAM17*^25^ for TNFR1 and *SLC34A1*^26^ for FGF-23 [Supplementary Table 2].

Clinical-stage targeting with any drug modality was reported for 35 of the 90 proteins on the Olink CVD-I panel [Supplementary Table 7]. Our MR analysis identified 11 proteins with causal evidence of involvement in human complex disease development that have not previously been targeted. Among those 11 proteins, four proteins were causal for a disease phenotype and did not show strong evidence of inverse causality with another phenotype (increasing specificity for intended indication), including CHI3L1 and SPON1 for atrial fibrillation and PAPPA for type-2 diabetes. Strong causal evidence was also identified for proteins targeted in phase-2 or later development. The MR evidence was concordant with drug indications for several protein targets but for some also suggested alternative indications or that monitoring of target-associated safety might be warranted. Monoclonal antibodies that block the CD40 ligand binding to CD40 – a critical element in T cell activation – have been shown to have positive clinical effects in patients with autoimmune diseases; but increased risk of thromboembolism precluded further clinical development^27^. These observations from clinical trials are in line with our findings that genetically *lower* levels of CD40 are associated with *lower* risk of RA, but *higher* risk of stroke. There are ongoing efforts to modify CD40L antibodies to retain efficacy while avoiding thromboembolism ^27^. However, our results suggest that decreasing circulating CD40 levels may have target-mediated beneficial effects on RA risk, while increasing the risk of ischemic stroke, i.e. that the increased risk of thromboembolism (manifest as stroke) is an on-target adverse effect. TRAIL-R2 is a key receptor for TRAIL, which has been shown to selectively drive tumour cells into apoptosis. Therefore, considerable effort to agonise TRAIL-R2 for treating cancers has been made in the past years^28^. We demonstrated that increased circulating TRAIL-R2 is protective against prostate cancer, which may suggest that this cancer type should be investigated in clinical trials evaluating the efficacy of TRAIL-R2 agonists.

Biomarkers are medical tools supporting clinical decision making and can be broadly classified as generic biomarkers for disease risk or prognosis, or as biomarkers reflecting the activity of specific disease processes or biology. Biomarkers that enable matching of target mechanisms to patient subgroups with greater than average benefit from treatment are enablers of precision medicine. Measuring biomarkers in a phase-II clinical trial of diabetic nephropathy, we show that CCR2/CCR5 small-molecule inhibition modulated circulating levels of CCL-4 and MCP-1, which may suggest that *trans*-pQTLs can guide selection of exploratory biomarkers to monitor the efficacy of specific target mechanisms. We also identified multiple complex traits causally affecting circulating protein levels. Such complex phenotype-to-protein associations may represent pathway-related causality to the complex phenotype of interest; or alternatively, ‘reverse causality’ which might pose an opportunity to evaluate implicated proteins as surrogate biomarkers for efficacy in interventional trials ^29^. For example, we found that higher BMI causally lowered RAGE, while higher circulating levels of RAGE were causally linked to a lower risk of T2D. Thus, developing a hypothetical therapeutic to increase RAGE (notwithstanding the other target-mediated effects discussed above) might represent a mechanism by which it is possible to off-set the risk of T2D arising from the global increases in obesity.

Protein-centric PRS’ may constitute an alternative approach to stratify individuals with genetic propensity for high circulating protein levels. While the genetic contribution to population variance was found to be higher for the CVD-I proteins than typically reported for complex traits, only 10 % of the protein-centric PRS’ explained 10 % or more of the protein variance in the independent replication cohort. One of those was ST2, a prognostic biomarker for heart failure^30^. ST2 showed evidence of inverse causality in asthma and positive causality in IBD. By constructing a genome-wide polygenetic risk score for ST2 levels from the MDC study, applying it to the UK Biobank and comparing asthma and IBD prevalence across eleven quantiles of the ST2 PRS, we validated the inverse causal role of ST2 in asthma and estimated the magnitude of ST2 increase required to decrease the risk of asthma to similar levels as individuals in the highest ST2 PRS category. Such use of PRS for proteins may be expanded to other disease endpoints and may be of use in precision medicine, to guide which patients may obtain most benefit from drugs that pharmacologically alter individual proteins.

In conclusion, our findings provide a comprehensive toolbox for evaluation and exploitation of therapeutic hypothesis and precision medicine approaches in complex disease. Such approaches provide an excellent opportunity to rejuvenate the drug development pipeline for new treatments.

---

**Supplementary Figure 1.** Chromosomal location of all primary associations at conventional GWAS significance

**Supplementary Figure 2.** Illustration of the online interactive tools for visualization of genomic loci, regions and plausible networks (www.scallop-consortium.com). **A.** Illustration of hotspot loci on chromosome 10 (left) and illustration of hotspot loci with independent effects established using COJO analysis (right) **B.** Circular Manhattan plot for TNF-R2. **C**. The pathway implicated by trans-pQTLs for plasma TNF-R2. The network shows the likely path from pQTL to TNF-R2.

**Supplementary Figure 3.** All proteins and tissues with significant PrediXcan correlations between protein and predicted mRNA levels.

**Supplementary Figure 4.** Effect of exposure to PF-04634817 on EN-RAGE, FGF-23, KIM-1, myoglobin and TNFR-2.

**Supplementary Figure 5.** Overview of protein levels having effect on complex phenotypes using Mendelian Randomization. Similar to figure 5B, but also showing effects with intermediate evidence strength.

**Supplementary Figure 6.** Overview of complex phenotypes having effect on protein levels using Mendelian Randomization.

**Supplementary Figure 7.** Decision tree describing the reasoning behind Mendelian Randomization evidence strength.

**Supplementary Figure 8.** Meta-regression of quantiles of ST2 polygenic risk score and relative risk of asthma (left) and inflammatory bowel disease (right). Values plotted on the x-axis relate to the quantile-specific mean difference in ST2 as compared to the 6th quantile. Values plotted on the y-axis relate to the quantile-specific log odds of disease as compared to the 6th quantile. The red line is the slope derived from the meta-regression across the ST2 quantiles of the PRS on log odds of disease, weighted by the standard error of the log odds.

**Supplementary Table 1.** Information about all measured proteins

**Supplementary Table 2.** List of all protein quantitative locus (pQTL) associations

**Supplementary Table 3.** Overview of protein-protein interaction (PPI) and text mining (TM) systems biology analysis

**Supplementary Table 4.** Systematic analysis of protein quantitative trait loci (pQTL) in previously published literature

**Supplementary Table 5.** Investigation of overlap between protein quantitative trait loci (pQTLs) and expression quantitative trait loci (eQTLs)

**Supplementary Table 6.** Summary-data-based Mendelian Randomization (SMR) using heterogeneity in dependent instruments (HEIDI) test.

**Supplementary Table 7.** Overview of gene products targeted by compounds or antibodies that have been in clinical development

**Supplementary Table 8.** Overview of participating cohorts

**Supplementary Table 9.** Overview of external genome-wide association study (GWAS) data used in mendelian randomization (MR) analyses

## Acknowledgments

MAK is supported by a Senior Research Fellowship from the National Health and Medical Research Council (NHMRC) of Australia (APP1158958). He also has a research grant from the Sigrid Juselius Foundation, Finland

Sources of Funding for SMCC, part of the national research infrastructure SIMPLER. We acknowledge the national research infrastructure SIMPLER (the Swedish Infrastructure for Medical Population-based Life-course and Environmental Research) for provisioning of facilities and support. SIMPLER receives funding through the Swedish Research Council under the grant no 2017-00644. This study was also supported by additional grants from the Swedish Research Council (grants no 2017-06100; no 2015-05997 and no 2015-03257), from the Swedish Research Council for Health, Working Life and Welfare (FORTE grant no 2017-00721) and Stiftelsen Olle Engkvist Byggmästare (grant no 2017/49)

Dr. Lubitz is supported by NIH grant 1R01HL139731 and American Heart Association 18SFRN34250007.

The Orkney Complex Disease Study (ORCADES) was supported by the Chief Scientist Office of the Scottish Government (CZB/4/276, CZB/4/710), a Royal Society URF to J.F.W., the MRC Human Genetics Unit quinquennial programme “QTL in Health and Disease”, Arthritis Research UK and the European Union framework program 6 EUROSPAN project (contract no. LSHG-CT-2006-018947). DNA extractions were performed at the Edinburgh Clinical Research Facility. We would like to acknowledge the invaluable contributions of the research nurses in Orkney, the administrative team in Edinburgh and the people of Orkney.

AB was supported by a Wellcome PhD training fellowship for clinicians (204979/Z/16/Z), the Edinburgh Clinical Academic Track (ECAT) programme

J. Gustav Smith and the genotyping of MPP-RES was supported by grants from the Swedish Heart-Lung Foundation (2016-0134 and 2016-0315), the Swedish Research Council (2017-02554), the European Research Council (ERC-STG-2015-679242), the Crafoord Foundation, Skåne University Hospital, the Scania county, governmental funding of clinical research within the Swedish National Health Service, a generous donation from the Knut and Alice Wallenberg foundation to the Wallenberg Center for Molecular Medicine in Lund, and funding from the Swedish Research Council (Linnaeus grant Dnr 349-2006-237, Strategic Research Area Exodiab Dnr 2009-1039) and Swedish Foundation for Strategic Research (Dnr IRC15-0067) to the Lund University Diabetes Center.

The study of the LifeLines-DEEP cohort is supported by the Netherlands Heart Foundation CVON grant 2018-27 to JF and AZ, Netherlands Organization for Scientific Research (NWO-Vidi grant 864.13.013 to JF, 016.178.056 to AZ, 917.14.374 to LF, Veni grant 194.006 to DZ, gravitation grant ExposomeNL to AZ, gravitation 024.003.001 to JF), European Research Council (ERC starting grant 715772 to AZ, 637640 to LF), LF also receives financial support from Oncode Institute.

We would like to thank Professor John Parks at Wake Forest School of Medicine, Winston-Salem, NC and Professor Daniel Rader at Perelman School of Medicine at the University of Pennsylvania, Philadelphia, PA for their kind donations of samples from transgenic mice and controls. This research has been conducted using the UK Biobank Resource under Application Number 13721.

## URLs

www.scallop-consortium.com

www.ebi.ac.uk/gwas/

www.proteinatlas.org

www.uniprot.org

https://www.pantherdb.org

david.ncifcrf.gov

clinicaltrials.gov

www.ebi.ac.uk/chembl

www.drugbank.ca

www.opentargets.org

neic.no/tryggve/

## Data availability

The full summary statistics of the Olink CVD-I protein GWAS have been deposited at the <u>SCALLOP-CVD-I online resource</u>, allowing access to interactive SCALLOP-CVD-I tools and unrestricted download access for secondary analyses. Additionally, a full copy has been deposited at https://doi.org/10.5281/zenodo.2615265 for long-term retention.

## Online Methods

### Cohorts and data collection

Summary statistics from GWAS of Olink CVD-I proteins were obtained from 13 cohorts of European ancestry. The details of all study cohorts are shown in [Supplementary Table 9]. Together the cohorts included a total of 21,758 individuals; although the average per-protein sample size was 17,747, since not all proteins passed quality control (QC) in all cohorts. Each cohort provided data imputed to 1000 Genomes Project phase 3 reference or later or to the Haplotype Reference Consortium (HRC) reference, which resulted in the testing of 21.4M SNPs. Because imputation schemes varied by cohort, this resulted in an average of 20.3M SNPs under investigation for each protein.

Each cohort applied quality control measures for call rate filters, sex mismatch, population outliers, heterozygosity and cryptic relatedness as documented in [Supplementary Table S08]. Prior to running the genetic analyses, NPX values of proteins (on the log_2_ scale) were rank-based inverse normal transformed and/or standardised to unit variance. Genetic analyses were conducted using additive model regressions, with adjustment for population structure and study-specific parameters [Supplementary Table 8]. Forest plots of cohort-specific effects are available for all significant and suggestive pQTLs using the <u>online tool</u>. Each contributing cohort uploaded the resulting summary statistics in a standardized format using a secure computational cluster provided by Neic Tryggve (https://neic.no/tryggve/). All meta-analysis was performed in duplicate at two different research centres using completely separate bioinformatic pipelines (L.F. and S.G.).

### Data cleaning and meta-analysis

A per-protein filtering threshold of >80% samples above the Olink detection limit was applied to each cohort, leaving data on 90 of the 92 proteins to be analysed. The remaining files had an average of 3% missing samples (per cohort statistics available in [Supplementary Table 8]). Minor allele frequencies were compared with those reported in 1000 Genomes EUR. A per-SNP filter was applied based on imputation quality level (at default setting for respective imputation algorithm) and minor allele count (at least 10 alleles per cohort). This resulted in the omission of 10% of the SNPs. Finally, meta-analysis was performed using METAL (2011-03-25) ^31^, applying the inverse-variance weighted approach (i.e. the STDERR option). *Cis*-pQTLs were defined as a signal within 1 Mb of the gene encoding the protein and all other signals were defined as *trans*-pQTLs.

### Replication analyses

We sought to replicate the findings in the Malmö Diet and Cancer (MDC) population-based cohort with 4,678 individuals, and in the Swedish Mammography Cohort Clinical (SMCC, part of the Swedish national research infrastructure SIMPLER described at www.simpler4health.se) population-based study of 4,495 women. In MDC, genotypes were imputed to the Haplotype Reference Consortium reference (HRC Unlimited v1.0.1) and data were analysed using linear regression in EPACTS 3.3.0 (linear Wald test). The genotypes in SMCC were measured using Illumina’s Global Screening Array and were imputed up to HRC v1.1 and 1000G phase3 (v5), and linear regressions of rank-based inverse-normal transformed protein values adjusting for age, storage time, and PC1-15 were performed using PLINK v2 (4 Mar 2019).

### Conditional and joint association analysis

To identify secondary signals at the 401 significant and suggestive loci identified, we performed analyses conditioning on the primary signal using conditional-joint analysis in GCTA (version 1.26.0)^32,33^. The Stanley cohort was chosen as an ancestrally well-matched LD-reference cohort. Meta-analysis summary data were processed with filtering for MAF (0.01) and r^2^ (<0.001) to ensure that secondary association signals identified were not driven by LD with the primary signal.

### Cross-reference of pQTLs with other complex traits

For each pQTL association, we searched PubMed and the EBI GWAS catalogue (URL: https://www.ebi.ac.uk/gwas/:November 2018) for published SNPs with any complex trait within 10kb or having an LD of r^2^>= 0.85.

### Comparison between eQTLs and pQTL

To identify eQTL that corresponded to each pQTL, we used three independent eQTL studies: LifeLines-DEEP ^34^, GTEx^35^ and eQTLGen^36^. Each SNP-protein pQTL pair was first converted to SNP-gene pairs using Olink platform protein identification and the gene annotation of Ensembl v91. Then, the significance of eQTLs for these SNP-gene pairs was assessed in three eQTL datasets, using two different cut-offs: a stringent genome-wide significance threshold (*P*<5×10^−8^) and a nominal significance of *P*<0.05.

In the eQTL dataset of LifeLines-DEEP, individual-level whole blood RNA-seq, protein and genotype data were available. This allowed for a direct comparison of the concordance of blood eQTLs and pQTLs. To do so, we re-tested eQTL associations for all pQTL pairs, using a previously published pipeline ^37^. The resulting eQTLs were considered genome-wide significant if it passed the permutation-based FDR <0.05 level, or to be nominally significant if the *P*-value was < 0.05.

In the eQTL datasets of GTEx v7 and eQTL-Gen, we did not have access to individual level data. Thus, the comparisons were conducted using publicly available eQTL results. In these datasets, we considered an eQTL genome-wide significant if it was within the reported genome-wide significant list, and nominally significant if it had a nominal *P*-value < 0.05. Altogether, if one pQTL pair had at least one significant eQTL effect in any dataset irrespective of allelic direction it was considered an overlapping pQTL-eQTL pair.

### Expression SMR analysis

We performed an SMR and HEIDI (heterogeneity in dependent instruments) analysis^8^ to identify the expression levels of genes that were associated with protein abundance through pleiotropy using pQTL summary statistics from this study and cis-eQTL summary data from published studies^38,39^.

The eQTL summary data used in the SMR analysis were from the Consortium for the Architecture of Gene Expression (CAGE), comprising 38,624 normalized gene expression probes and ~8 million SNPs from 2,765 blood samples. The eQTL effects were in standard deviation (SD) units of expression levels. We excluded the gene probes in the major histocompatibility complex (MHC) region and included only the gene probes with at least one cis-eQTL at *P*<5×10^−8^ (a basic assumption of SMR), resulting in 9,538 gene expression probes.

The SMR test uses a SNP instrument (i.e., the top associated eQTL) to detect association between two phenotypes (i.e., gene and protein in this case). The HEIDI test utilises LD between the SNP instrument and other SNPs in the cis-region to distinguish whether the association identified by the SMR test is driven by a set of shared genetic variants between two traits (pleiotropic or causal model) or distinct sets of variants in LD (linkage model)^8^. Only the associations that surpassed the genome-wide significance level of the SMR test (*P*_SMR_ < 0.05 / *m* with *m* being the number of SMR tests) and were not rejected by the HEIDI test (*P*_HEIDI_ > 0.01) were reported as significant.

### PrediXcan and transcript-wide association of CVD-I protein levels

Imputation of gene expression was performed in the IMPROVE study. After standard quality control, genotypes were pre-phased using Eagle2, and then subsequently imputed by minimac4 using the 1000 Genomes reference. A filter on RSQ 0.8 and minor allele frequency 0.01 was set on the imputed genotypes prior to prediction with PrediXcan, which used 44 tissue models based on GTEx v7.

Using protein data collected on the CVD-I chip in the same individuals, the associations between protein levels in plasma and the predicted expression of their respective coding gene across 20 tissues (from the PrediXcan model) were modelled by a linear model in R. False discovery rate were estimated based on Q-values (using the R package qvalue). In total, 64 genes in one to 18 tissues were tested for associations between protein levels and predicted expression. Heatmaps were constructed (using the pheatmap package in R) for any gene with a significant association (FDR<0.05) in at least one tissue.

### Systems Biology

Two sets of network analysis were performed, one using the protein-protein interaction (PPI) data from the inBio Map™ (InWeb_InBioMap) and one using significant associations from text-mining (TM). These two networks each had 13,033 and 14,635 nodes, respectively; and 147,882 and 193,777 edges, respectively. In both setups, the shortest path between any of the cis-gene intermediaries to the protein was identified; altogether 10,222 pairs were compared. Of the 372 trans-pQTL associations reported in [Supplementary Table S02], 335 associations had both cis-gene intermediaries and plasma protein in the network allowing their analysis. The likelihood of a path arising by chance was calculated by permutation sampling, using 1,000,000 random networks were generated with a conserved degree distribution. A new algorithm was developed for *de novo* random network generation, which generated random networks with a nearly conserved degree distribution in a feasible time-frame. Further details are available in [Supplementary Notes 1].

### Assignment of cis-intermediary genes

To assign the most plausible causal gene for each of the CVD-I trans-pQTLs we applied a hierarchical approach based on analysis of InWeb_InBioMap PPI, TM, and genomic distance between gene and lead variant at each locus. Results were then manually reviewed by literature, gene expression analysis (proteinatlas.org) and published pQTLs which led to the re-assignment of 52 genes.

### Human in-vivo validation of trans-pQTLs

PF-04634817 is a competitive dual inhibitor of CCR2 and CCR5 receptors. In the recent B1261007 study, (ClinicalTrials.gov Identifier: NCT01712061), samples were collected from subjects with diabetic nephropathy and treated with PF-04634817 for 12 weeks. CCL-2 (MCP-1) was measured in serum by ELISA at Eurofins (The Netherlands). CCL4 (MIP-1b) and CCL-8 were measured in plasma using Luminex assays (Bio-Rad, Berkeley, CA). CCL5 (RANTES), was measured in plasma as part of a multi-analyte panel at Myriad Rules Based Medicine (Austin, TX).

### Mouse in-vivo validation of trans-pQTLs

Plasma from transgenic- and matched control mice were randomised on a PCR plate. The samples included five mice with targeted deletion of hepatocyte ABCA1^17^ together with five matched control mice, three mice with whole-body TRIB1^18^ knockdown and three controls and four mice with liver-specific knockdown of TRIB1 and four matched controls. Protein levels of stem cell factor (SCF) was measured using the Olink PEA Mouse exploratory panel according to the manufacturer’s instruction (Olink Proteomics, Uppsala, Sweden). The plasma levels of SCF were normalised against average protein concentrations using information on an additional 91 proteins. TRIB1 whole-body and liver-specific mice were analysed jointly as were the respective wild-type controls. The median plasma levels of SCF were compared using the Mann-Whitney U test for unpaired samples.

### Mendelian Randomization

To study the causal effects of the protein on selected disease outcomes, we performed two-sample Mendelian randomization analyses. We created two sets of instrumental variables (IVs) for each of the 85 proteins with variants reaching multiple testing-corrected significance in our discovery GWAS: (a) *cis* IVs including one or more independent variants (LD r^2^=0.001 within ±1Mb of the transcript boundaries of the gene encoding the protein); and (b) *pan* IVs including all independent (LD r^2^=0) variants associated with the protein, i.e. combining *cis* and *trans* pQTLs. The per-allelic beta coefficients from the main GWAS analyses were used as weights in the IVs. For the outcomes, we obtained the relevant SNP-to-trait summary statistics from publicly-available GWAS as outcomes [Supplementary Table 9]. When lead variants from our main GWAS were not available in these summary statistics, we replaced them with proxies (LD r^2^>0.85). For each individual SNP-protein and SNP-outcome association, we generated an instrumental variable Wald ratio estimate, with standard errors obtained using the delta method. When the instrument included more than one SNP, summary IV estimates were generated by combining individual SNP Wald estimates by inverse-variance weighted fixed-effect meta-analysis. We report associations with a Benjamini-Hochberg false discovery rate (FDR) ≤ 5%, applied separately to summary estimates from *cis*-pQTL and *pan*-pQTL IVs, using pooled estimates for all four diseases. We graded the evidence of causality using a framework outlined in [Supplementary Figure 7], using the following categories: <u>strong</u> (*cis*-IV estimate FDR≤ 5%); <u>intermediate</u> (pan-IV estimate FDR≤ 5% with: (i) no heterogeneity between cis-IV estimate and pan-IV estimate; *and* (ii) no evidence of the MR estimate being unduly influenced by a *trans*-pQTL in leave-one-out analysis); or <u>weak</u> (pan-IV estimate FDR≤ 5% but: no *cis*-pQTL IV available; heterogeneity between *cis*- and all-IVs; or evidence of undue influence by a trans-pQTL). Heterogeneity between *pan*-IV and *cis*-IV estimates were calculated using Cochran’s Q tests, with *P*<0.05 denoting evidence against the null hypothesis, and applying a Bonferroni adjustment for multiple testing. Mendelian randomization was conducted in duplicate by two separate analysts and analyses were performed in Stata (StataCorp, Texas, USA) version 13.3 using the *mrivests*, *metan* and *multproc* commands and R. Of the 2437 IV estimates derived using *cis*-pQTL instruments across the 85 proteins and 38 outcome traits, the IV estimates of 50 protein-to-disease associations met the FDR≤5% (corresponding to an uncorrected *P*≤1.1×10^−3^). Of the 3044 IV estimates composed using all pQTL instruments, 281 IV estimates met FDR≤ 5% (corresponding to *P*≤ 4.7×10^−3^; [Figure 5A]. The decision tree for scoring the strength of MR evidence is available in [Supplementary Figure 7].

### Heritability analyses

We estimated the total SNP-heritability (h_SNP_^2^) for the plasma level of each protein from the summary statistics of each individual GWAS by summing the contributions from two independent partitions of the SNPs: primary major loci and polygenic background. We defined the variance explained by primary major loci (major loci h_SNP_^2^) as the sum of the estimated variance explained (2*β^2^,*f*(1-f)), where f is the minor allele frequency, and owing to the fact that the phenotypic variance has been standardized across lead SNPs indexing all primary genome-wide significant loci. We used LDSC regression^40^ to estimate the contribution of the polygenic background (polygenic h_SNP_^2^) for each protein, which we define as the contribution of all loci not indexed by a genome-wide significant lead SNP. LDSC regression is known to perform poorly when large effect, major genes are present, as it was derived under the assumption of a simple polygenic genetic architecture^40^. To account for this and avoid double counting the variance explained by major loci through LD surrogates, prior to estimating the LDSC regression polygenic h_SNP_^2^, we censored all SNPs within 10 Mb of genome-wide significant lead SNPs for all primary loci.

### Polygenic risk score calculation

Polygenic risk scores were derived using LDpred algorithm^41^, which adjusts the effect of each SNP allele for those of other SNP alleles in linkage disequilibrium (LD) with it, and also takes into account the likelihood of a given allele to have a true effect according to a user-defined parameter, which we used as all 7 default LDpred-settings, with values from 1 through 1×10^−5^. The algorithm was directed to use HapMap3 SNPs that had a minor allele frequency >0.05, Hardy-Weinberg equilibrium P>1e-05 and genotype-yield >0.95, consistent. Variance explained in the independent MDC-study was tested according to a step-wise model, first including non-genetic covariates, then additional variability explained by adding SNPs from genome-wide significant SNPs (major loci V.E._PRS_), and then additional variability explained by adding the 7 LDpred-derived scores as additional covariates (polygenic V.E._PRS_).

### ST2 polygenic risk score for asthma and inflammatory bowel disease in the UK biobank

Prior to analysis subjects who were not White British (based on self-reported ancestry in combination with genetic PCA) in the maximum unrelated subset were filtered out. All bi-allelic SNPs with MAF >= 1% and MaCH rsq >= 0.8 were kept. The Z-score transformed LDpred PRS (wt2) for ST2 was calculated as described for MDC in 337,484 White British UK Biobank participants. Association with asthma and IBD were tested using logistic regression adjusting for age, sex, PC1-10, genotype batch using either the continuous PRS or the PRS quantile-bins as predictors. The UK Biobank protocol has been described previously^42^ and is available online (https://www.ukbiobank.ac.uk). The genotype quality control (QC), phasing, and imputation was performed centrally and has been previously described ^43^. Outcomes (defined based on self-reported data at baseline and/or the inpatient and death registry [including primary and secondary causes as well as prevalent and incident disease]) Asthma: Self-reported touchscreen (6152), self-reported nurse interview (20002), or ICD-10 “J45”. Conflicting self-reported results set to missing unless “J45” was reported. Inflammatory bowel disease: nurse interview (20002) or ICD-10 K50-K52.

### Meta-regression analysis for ST2 PRS, asthma and IBD

We estimated the per-quantile and per-SD associations of the weighted PRS for ST2 (MDC study) on risks of asthma and IBD (UK Biobank) by taking the quantile associations with ST2, asthma and IBD and conducting meta-regression analyses whereby the dependent variable was the quantile-specific logOR and corresponding SE of asthma or IBD and the independent variable was the quantile specific beta coeffient for ST2. This was conducted using the “metareg” package in STATA SE v13.1 (Statacorp, USA). Plots from the metaregression are presented in [Supplementary Figure 8].

