## Supplementary Notes 1 for "Genomic evaluation of circulating proteins for drug target characterisation and precision medicine"

**Systems biology**

The purpose of the network analysis was to identify the most likely chain of events (via protein interactions or other forms of regulation) leading from the trans-pQTL to the plasma protein being affected. For each of the trans-pQTL signals, all genes within a distance of 1 Mb of the SNP were considered to be a possible cis-gene intermediary between the SNP and the plasma protein. For investigating possible chains of events acting via physical interactions, we utilized the protein-protein interaction (PPI) network inBio Map™ (InWeb_InBioMap)^1^

(<https://www.intomics.com/inbio/map.html>), and for investing other potential regulatory chains we generated an association network based on text mining of PubMed abstracts. The inBio Map PPI resource was selected because it is built entirely on experimentally determined interactions, has a large coverage of the human interactome and furthermore, provides a confidence score on the experimental support for each edge in the network allowing for fine-grained control of what to include. In this study, we only used the high confidence subset of the global PPI network (see details below). Possible chains of events were then defined as paths in the PPI network connecting the protein product of the possible cis-gene intermediaries to the plasma protein whose protein level was affected. Under the assumption that the shortest path is also the most likely path, we calculated the statistical significance of each path by a network permutation scheme where the shortest path of each cis-gene intermediary was compared to the shortest path in a set of randomized networks.

For this approach to work, it is important that the random networks are similar to the original PPI network. Hence, they should contain the same number of nodes (proteins) and edges (interactions) as inBio Map. Furthermore, it has been shown that biological networks belong to a class of networks called scale-free networks and that their node degree distribution follow a power-law^2^. Therefore, it is important that the random networks also display this property and in particular, that they follow the same node degree distribution as inBio Map.

Random networks can be generated either by re-wiring the existing network (by pairwise edge swapping) or by building a random network from scratch (*de novo* network generation). However, with the requirement of a conserved node degree distribution, these methods become prohibitively slow (generating 1,000,000 random networks would take months-years even on a multicore supercomputer). Previous approaches to randomization of the complete protein-protein interactome have overcome this limitation by re-wiring the existing network but only swapping edges between certain subsets of nodes in the network (e.g. nodes with the same node degree)^3^.

We developed an algorithm that creates *de novo* random networks with a nearly (99.97 %) conserved node degree distribution of inBio Map. Since the networks are randomly generated from scratch, they will be completely independent of the original PPI network but they will still maintain the required properties.

Our algorithm exploits the fact that a random network with a conserved node degree distribution can be generated relatively fast, if the network is allowed to contain multi-edges and self-edges (loops). The multi-edges and loops are subsequently removed and the lack of edges is accounted for by generating a new network filling in the missing edges. This procedure is repeated a number of times until the random network is sufficiently similar to the original network.

The algorithm was implemented using the igraph package^4^ for Python (<https://igraph.org/python/>) and is outlined below:

**Iterative refinement of *de novo* generated random networks:**

1. Calculate the node degree distribution of the original network (*N_O_*)
2. Generate a random network (*N_A_*) with the same node degree distribution as *N_O_*
3. Remove multi-edges and loops from *N_A_*
4. Iterate until the stop criterion is met:
   a. Calculate the difference between the node degree distribution of the random network (*N_A_*) and the original network (*N_O_*)
   b. Generate a new random network (*N_B_*) with node degree distribution equal to the difference between *N_A_* and *N_O_* just calculated
   c. Merge the two random networks *N_A_* and *N_B_*
   d. Remove multi-edges and loops from the merged network – this is now the new random network (*N_A_*)

After each iteration of the algorithm, the distance between the original network (*N_O_*) and the random network (*N_A_*) was evaluated. The distance was defined as the sum of all pairwise absolute differences in node degree between all nodes (*n*) in the original network (*N_O_*) and the random network (*N_A_*):

$$distance=\sum_{n} | degree(n_{A})-degree(n_{O}) |$$

The stop criterion was defined as “the distance being less than or equal to 100” or if “the distance did not change for 5 consecutive iterations”.

The algorithm was applied to the high confidence subset of the PPI network inBio Map (version 2018_04_05), defined by the edges with a confidence score >= 0.117. Unspecific post-translational modifications (PTMs), such as the attachment of ubiquitin to target proteins, are often picked up by the methods used to detect protein-protein interactions. To avoid a bias from PTMs in our analysis a few proteins often associated with PTMs were left out from the analysis (UBB, UBC, UBA52, RPS27A, SUMO1-4). The final PPI network used in the analysis contained 13,033 proteins (nodes) and 147,882 high confidence interactions (edges). The total permutation time, generating 1,000,000 random networks using 20 CPU cores, was 6.7 hours.

The average distance (as defined above) between the random networks and inBio Map was 83.1 with a standard deviation of 10.6. This means that the node degree distribution of the random networks was highly conserved and on average 99.97% identical to the node degree distribution of inBio Map.

**Significance of shortest paths**

The purpose of generating random networks was to evaluate the significance of the shortest paths between the cis-gene intermediaries and the plasma proteins for each of the trans-pQTL signals. Thus, for each potential cis-gene intermediary and the plasma protein, the shortest path was also calculated in each of the 1,000,000 random networks and the significance of the observed shortest path was then calculated as the fraction of random networks with a shortest path, shorter than or equal to the observed shortest path in inBio Map. As multiple paths were tested for each trans-pQTL signal, the Benjamini–Hochberg procedure was used to correct for multiple testing, yielding FDR controlled q-values.

Calculation of the shortest paths was actually more time-consuming than the permutation of the network, and the running time of the whole PPI network analysis was 37.3 hours using 28 CPU cores.

The results of the PPI network analysis are available in Supplementary Table S08.

**Text mining association network**

The inBio Map PPI network consists of experimentally observed physical interactions between proteins. Some interactions between proteins (e.g. protein phosphorylation by a kinase) are very short-lived and may not be picked up by some of the methods (especially co-purification based methods) used to detect physical interactions between proteins but they might still be relevant. Some of these interactions are described in the scientific literature and can be picked up by text mining.

Therefore, as an alternative to the PPI network, we also created a network of statistically significant associations between proteins identified by text mining (TM network). We used the text mining resource inBio Know™ (<https://www.intomics.com/technology/#text-mining>) to identify proteins co-mentioned in PubMed abstracts. inBio Know is based on highly curated protein synonyms for the human proteome and provides a statistical framework to identify proteins that are significantly associated in the scientific literature. Briefly, based on the manually curated protein synonyms, the occurrence of each protein from Uniprot was determined in PubMed abstracts (PubMed download date: 2018_05_03). Only human proteins with the status “reviewed” in Uniprot were considered. The pairwise co-occurrence between proteins was determined and Fisher’s exact test was used to calculate the significance of the association. A conservative, multiple-testing corrected *p*-value cutoff of 10^-8^ was applied to obtain significant associations between proteins.

A TM network was then created by connecting proteins that were significantly associated in PubMed abstracts. The TM network contained 14,635 proteins (nodes) and 193,777 associations (edges).

The algorithm described for the PPI network above was applied to the TM network in order to find significant paths between the potential cis-gene intermediaries and the plasma proteins in the text mining associations. Using the same stop criterion as above the total running time for the analysis was 87.9 hours using 28 CPU cores. Due to the topology of the TM network, the algorithm did not converge as fast as for the PPI network. This was also reflected in the average distance of the random networks, which was 739.0 with a standard deviation of 41.7. The distance could be reduced at the expense of longer running times but since the degree distribution of the random networks was still 99.81% identical to the degree distribution of the original TM network, we did not find that necessary.

Finally, we created a combined PPI and TM network consisting of both the protein-protein interactions and the associations identified by text mining simply by merging the two networks (PPI+TM network). This network contained 16,150 proteins (nodes) and 314,503 interactions (edges). The running time of the network analysis was 126.9 hours using 28 CPU cores. The average distance of the random networks was 370.3 with a standard deviation of 30.5. The degree distribution of the random networks was 99.94% identical with the degree distribution of the original PPI+TM network.

The results of the TM network analysis and the PPI+TM network analysis are both available in Supplementary Table S08.
